# Does the choice of nucleotide substitution models matter topologically?

**DOI:** 10.1101/041566

**Authors:** Michael Hoff, Stefan Orf, Benedikt Riehm, Diego Darriba, Alexandros Stamatakis

**Affiliations:** Karlsruhe Institute of Technology, Department of Informatics, Kaiserstraße 12, 76131, Karlsruhe, Germany; The Exelixis Lab, Scientific Computing Group, Heidelberg Institute for Theoretical Studies, Schloss-Wolfsbrunnenweg 35, 69118 Heidelberg, Germany

**Keywords:** phylogenetics, nucleotide substitution, model selection, information criterion, BIC, AIC

## Abstract

**Background:** In the context of a master level programming practical at the computer science department of the Karlsruhe Institute of Technology, we developed and make available an open-source code for testing all 203 possible nucleotide substitution models in the Maximum Likelihood (ML) setting under the common Akaike, corrected Akaike, and Bayesian information criteria. We address the question if model selection matters topologically, that is, if conducting ML inferences under the optimal, instead of a standard General Time Reversible model, yields different tree topologies. We also assess, to which degree models selected and trees inferred under the three standard criteria (AIC, AICc, BIC) differ. Finally, we assess if the definition of the sample size (#sites versus #sites × #taxa) yields different models and, as a consequence, different tree topologies.

**Results:** We find that, all three factors (by order of impact: nucleotide model selection, information criterion used, sample size definition) can yield topologically substantially different final tree topologies (topological difference exceeding 10%) for approximately 5% of the tree inferences conducted on the 39 empirical datasets used in our study.

**Conclusions:** We find that, using the best-fit nucleotide substitution model may change the final ML tree topology compared to an inference under a default GTR model. The effect is less pronounced when comparing distinct information criteria. Nonetheless, in some cases we did obtain substantial topological differences.

## Background

Statistical models of DNA evolution as used in Bayesian inference (BI) and Maximum Likelihood (ML) methods for phylogenetic reconstruction are typically required to be time-reversible. That is, evolution (or the Markov process modeling it) is assumed to occur in the same way if followed forward or backward in time. Time-reversibility has some intrinsic computational advantages, mainly that, the likelihood of a tree one intends to score is the same for any placement of the virtual root that is deployed to direct the Felsenstein pruning algorithm [1].

To ensure that, a nucleotide substitution matrix is time-reversible, it must exhibit a certain symmetry. This symmetry requirement is depicted in the following example for a set of substitution rates {*r*_AC_, *r*_AG_, *r*_AT_, *r*_CG_, *r*_CT_, *r*_GT_} and a set of stationary frequencies {*π_A_*, *π_C_*, *π_G_*, *π_T_*}:

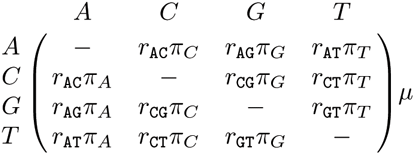

As implied by the above representation the rates r_XY_ in the upper and lower diagonal part of this matrix must be symmetrical. Therefore, there can be at most 6 distinct, independent rates. Because rates in such a matrix are relative rates, one rate (typically *r*_GT_) is set to 1.0 by default such that there are at most 5 free parameters that can be estimated via ML optimization or sampled with MCMC methods for BI. A time-reversible DNA substitution matrix can have between 1 up to 6 distinct rates. Note that, we further differentiate between substitution models with identical base frequencies (i.e., *π_A_* = …*π_t_* = 0.25) or distinct base frequencies. Distinct base frequencies can be either be obtained by using empirical base frequencies or via direct sampling (BI) or ML inference of the base frequencies. Nonetheless, here we mainly focus on models as defined by substitution rates and not base frequencies. If all 6 rates and all base frequencies are identical we obtain the Jukes-Cantor (JC [2]) model and if all 6 rates are independent, we obtain the General Time-Reversible (GTR [3]) model.

Evidently, there is a large number of possible time-reversible models situated between the JC and GTR model. We can essentially consider all possible combinations of dependencies between the 6 substitution rates. In the literature, only a subset of these has been explicitly described and named, for instance the HKY85 model [4]. We are aware of two papers and respective implementations that consider all possible time-reversible nucleotide models and intend to select the best-fitting one among these 203 substitution matrices.

The first paper that addressed the problem used Bayesian statistics and MCMC methods to sample among all possible substitution matrices proportional to their posterior probability [5]. The method for integrating over all substitution models is available in MrBayes version 3.2 [6] via the command `lset nst=mixed`.

Another code that allows for testing all possible substitution matrices in a ML framework based on the AIC (Akaike Information Criterion), AICc (corrected AIC) BIC (Bayesian Information Criterion), and DT (Decision Theory) criteria is `jModelTest 2` [7]. However, `jModelTest` relies on `PHYML` [8] to calculate likelihood scores for a given substitution model and tree. `PHYML` is relatively slow compared to `RaxML` (v. 8.1.7 [9]) and the Phylogenetic Likelihood Library (PLL [10]) that relies on the same phylogenetic likelihood kernel implementation as `RaxML`. For instance, optimizing the model parameters and branch lengths on a fixed tree with 354 ITS sequences [11] using a GTR model with Γ-distributed rate heterogeneity among sites [12] and 4 discrete rate categories requires 11 seconds with `RaxML` and 110 seconds with `PHYML`.

Here, we present and make available a fast implementation of a ML-based model testing tool for all possible time-reversible DNA substitution models. It was developed by three students within the context of a programming practical at the Master level in the computer science department of the Karlsruhe Institute of Technology. The educational aspects of this project are discussed in the on-line supplement.

The effect of different selection criteria and substitution models on ML-based phylogenetic analyses has already been discussed before. Darriba *et al.* [7] evaluated the accuracy of distinct model selection criteria with respect to recovering the *true* generating model parameters (e.g., substitution rates, proportion of invariant sites, or the α shape parameter of the Γ distribution) under simulation, using 40, 000 synthetic alignments under a wide range of model parameters.

Ripplinger and Sullivan [13] evaluated the influence of model choice on empirical tree topologies as determined by using the hierarchical Likelihood Ratio Test (hLRT [14]), the AICc and BIC criteria, as well as DT. In their tests, the selection criteria at hand, often returned different best-fit models. These different best-fit models then also induced distinct ML trees. Ripplinger and Sullivan exclusively focused on selecting named models in their study, that is, they did not select among all possible 203 substitution models. They did, however, also investigate the impact of model selection on bootstrap support values. Note that, this was not feasible in our study because of the limited amount of time and credits available to the students for completing the programming practical.

Nonetheless, our work fills the gap by the two aforementioned papers [7, 13] since we study the impact of selecting among all possible 203 substitution models on ML-based tree inferences on empirical datasets compared to tree inferences conducted under a default GTR+Γ model.

Initially, we present two algorithms for generating all possible substitution models. Thereafter, we discuss the sequential and parallel implementation using the PLL. subsequently, we present a set of experiments on empirical datasets to answer the following question: Does model selection really matter with respect to its impact on the shape of the final tree topology? Posada and Buckley discussed the potential impact of the sample size on AICc and BIC criteria [15]. We also assess if different possible definitions of the sample size as used for calculating information criteria influence the model that will be selected. In addition, we conduct a small simulation study to verify that our tool works correctly. We conclude with speedup data for the parallel implementation of our tool.

## Algorithms

The model test needs to calculate the ML score for every possible time-reversible nucleotide substitution model. A full list of all possible models was already available [5]. Nonetheless, part of the programming task was to design algorithms for generating them.

In the following, we first provide some background information and notation and then describe our two algorithms for generating all time-reversible nucleotide substitution models.

### Notation, Terminology & Properties

In analogy to [5] we encode a substitution model as a string of six digits over Σ_6_:= {1, 2, 3, 4, 5, 6}. For example the string `111333` describes a substitution model with the following constraints: *r*_AC_ = *r*_AG_ = *r*_AT_ and *r*_CG_ = *r*_CT_ = *r*_GT_. Note that, the PLL [10] offers a similar format for specifying time-reversible nucleotide substitution models. The only difference is that it uses the alphabet {0, 1, 2, 3, 4, 5}.

In this string notation, the GTR model is denoted by 123456, that is, all rates are different *and* independent from each other. The HKY85 model is denoted by `121121`. Here, we constrain *r*_AC_ = *r*_AT_ = *r*_CG_ = *r*_GT_ and *r*_AG_ = *r*_CT_.

We call a model a *k–model*, if it has exactly *k* distinct substitution rates. Hence, the parameter *k* characterizes the degree of freedom of a model. We further define *Σ*_*k*_ := {1, 2,…,*k*} as the alphabet of all digits with value ≤ *k*.

While two distinct models over *Σ*_*6*_ can consist of different digits/characters, they can specify *exactly* the same constraints. For example, `121121` and `212212` are semantically identical since they have the same degree of freedom and identical rate dependencies. We call such models *redundant.*

To avoid generating redundant models we introduce a *normalization property*, which holds only for one, unique model in a set of semantically identical models. A model string *s =* s_1_s_2_s_3_s_4_s_5_s_6_ (e.g., `123456` induces s_1_ = 1, s_2_ = 2,…) with digits s_*i*_ ∈ Σ_6_ is *normalized, if and only if*, for each *first* occurrence of a value s_*i*_ in *s* the following holds: {s_1_, s_2_,…, s_*i*−1_} = Σ_s_*i*_−1_. For example, the first occurrence of digit 3 in a model requires the prefix, excluding the current digit, to consist of exactly two digits: 1 and 2. The models `123111, 122132, 123456` fulfill this constraint and the models `311111, 113111, 124356` do not. The semantically equivalent, normalized models `122222, 112111, 123456` for the unnormalized ones in our example can be obtained by consistently replacing characters from left to right. As a consequence of the above definition, a normalized *k–*model therefore only comprises characters in *Σ*_*k*_.

We further define the *k–prefix* of a *k–*model *m*, as the shortest prefix of *m* that contains all characters in *Σ*_*k*_ at least once. This part of the string is denoted by underlining the respective characters. For example, the 4–model <u>1243</u>11 has the 4–prefix 1243, as any shorter prefix, say 124, does not yield all characters in Σ_4_ = {1, 2, 3, 4}.

In the following, we first present a brute-force algorithm and then a more elegant constructive algorithm for generating all model strings.

### Brute Force Approach

Our brute-force implementation initially enumerates all possible strings of length 6 for the alphabet Σ_6_ = {1,2,3,4,5,6}. The enumeration is done by counting from `111111` to `666666` in the base-6 numerical system, except that the digits are in the range 1… 6 instead of 0 … 5. In total, this yields 6^6^ = 46656 strings out of which only 203 are non-redundant.

To ensure that the algorithm only returns normalized model strings, we normalize every enumerated string and subsequently filter out redundant models.

The normalization re-maps the digits of the model string such that the normalization property holds. In the model string, every constraint is represented by a unique digit. The transformation now chooses new digits for each constraint. These digits are assigned incrementally starting with 1. For this, the algorithm identifies the constraints of the model string by traversing it from left to right. To properly order the constraints, they are ordered by the index of their corresponding leftmost digit.

The first occurrence of each normalized string encountered in the enumeration of all strings is then added to a final model list. We discard normalized model strings that have already been included in this list. When all strings have been enumerated, normalized, and added to the final model list, the list contains all 203 non-redundant model strings.

Note that, a normalized string always begins with 1, the second digit can only be 1 or 2 etc. We use this observation to optimize the brute force enumeration algorithm. For this, we only place characters in the set {1,…, *n*} at the nth position of the string. This prevents the algorithm from enumerating a large number of non-normalized strings. With this modification, only 1*2*3*4*5*6 = 720 strings are enumerated. This generates only 1.5% of the strings the naive brute-force approach enumerates and yields only 720 – 203 = 517 redundant models.

### Inductive Algorithm

Our inductive algorithm uses the set of all *k–*models to construct all possible (*k* + 1)–models. The algorithm starts with the single 1–model `111111` (the JC model) and constructs all 31 normalized 2-models (e.g., 111112, 111121, 111122, …). This process continues inductively by incrementing *k* until the sole possible 6-model 123456 (the GTR model) is generated.

In the following we will explain how a single new (*k* + 1)–model can be derived by applying *k* ↦ *k* + 1 on a *k*-model by using character replacement. Extending a normalized model to *k*+1 degrees of freedom requires one character to be replaced by a new character *k* + 1. To maintain the normalization property and to ensure the correctness of this operation, the replacement must not be applied to the *k*-prefix. For example when applying 2 ↦ 3 the 2-prefix 12 of model <u>12</u>1111 must not be used for a character replacement operation, since the resulting models 321111 and 131111 would not be normalized. To avoid the construction of redundant models, we further restrict the character replacement step to only replace characters of value *k*. Otherwise, the sets of (*k*+1)–models that are derived from different *k*-source models will not be disjoint. To illustrate what can happen, the step 2 ↦ 3 is applied to the third character of two 2-models <u>12</u>1111 and <u>12</u>2111. This will generate the 3-model <u>123</u>111 twice. An example for a derivation respecting both constraints is <u>123</u>3211 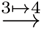 <u>1234</u>211. Note that, the (*k* +1)–prefix always extends the *k*-prefix to include the newly inserted character *k* + 1.

The algorithm relies on the property that every conducted replacement operation yields a valid model and thus, a fraction of the overall result. As a consequence, every model derivation step has to be applied to a copy of the source model to avoid over-writing already generated models.

Given a single *k*-model, the induction step *k* ↦ *k* + 1 is conducted in two phases. First, all characters not contained in the *k*-prefix that have value *k* are used to derive new models, one for each such character. This generates a set of (*k* + 1)–models, where the (*k* + 1)–prefixes now protect their newly inserted characters. For example the 1-model <u>1</u>11111 produces the 2-models <u>12</u>1111, <u>112</u>111, <u>1112</u>11, <u>11112</u>1 and <u>111112</u>. Note that, only the first phase derives (*k* + 1)–models from *k*-models and thus produces models with a longer protected prefix.

In the second phase, we use this set of (*k*+1)–models to recursively derive further (*k* + 1)–models. For each model in the set, we identify all occurrences of *k* not contained in the (*k*+1)–prefix. For each such “unprotected” occurrence of *k* we generate a new model by replacing the character *k* by *k* + 1. Each new model is then processed recursively in the same way. Note that, only occurrences of *k* right of the last replaced character are considered. This process continues until we reach the end of the string.

For example the model <u>1112</u>11, which has been generated in the first phase of 1 ↦ 2 yields 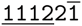 and <u>1112</u>12. Notice the changes in both new models and the trailing 1 in the first model marked by a line on top of it. The line indicates that the recursive processing of this model will start at this character and not consider characters left of it. The recursion then yields <u>1112</u>22, as the marked 1 can still be replaced.

Figure 1 shows a more complex derivation process. The 2-model 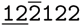 is used to derive 3–models starting at the left-most digit not contained in the prefix. Edges represent both, continuation of iteration and start of recursion on newly created models. Each first replacement increases the degree of freedom by one. Hence, the prefix is extended and the second phase of processing begins on this path.

**Figure 1.**
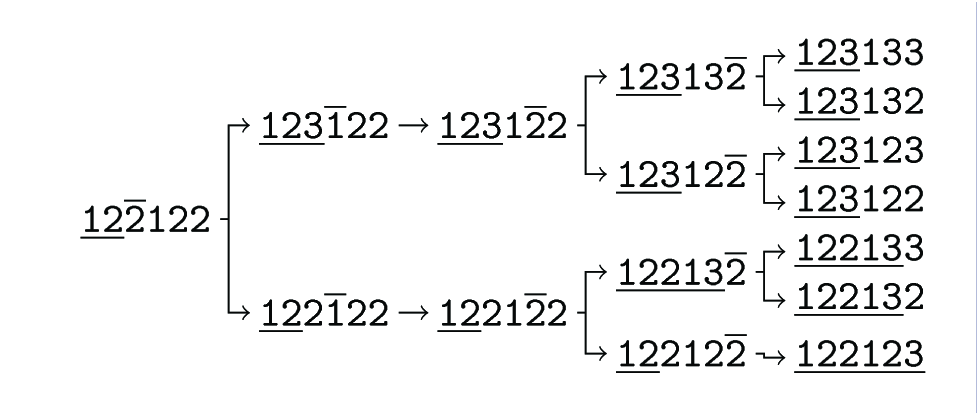
Inductive model derivation step 2↦3. Process tree showing how all 3–models derivable from 122122 are generated. Lines on top of characters indicate execution progress. Lines beneath visualize the 2– respectively 3–prefix.

Given a set of *k*-models, the induction step *k* ↦*k* + 1 as explained above is applied to each model in the set. The hereby obtained sets of (*k*+1)–models are disjoint by construction and can thus be easily unified to one complete set of (*k* + 1)–models. The overall induction process can be visualized by the following sequence of inductive steps:
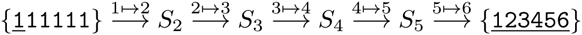

where |*S*_2_| = 31, |*S*_3_| = 90, |*S*_4_| = 65 and |*S*_5_| = 15.

## Implementation

The goal of the programming practical was to develop a program that tests all possible time-reversible nucleotide substitution models under Maximum Likelihood (ML) on a fixed, reasonable tree and subsequently selects the best model. Model selection is conducted using the following standard criteria: AIC, BIC, and AICc. In other words, our algorithm represents an ML-based re-implementation of the Bayesian approach deployed in [5]. Once the best model has been selected, the tool shall also conduct a ML-based phylogenetic tree search under the best model.

For likelihood calculations, tree searches, and handling I/O we used PLL [10].

Initially, we provide a top-level outline of the algorithm:

- Parse a multiple sequence alignment of DNA sequences
- Compute a reasonable, non-random, reference tree using parsimony
- Generate all 203 possible time-reversible substitution matrices
- Loop over these 203 matrices and optimize the ML score on the fixed tree for each model using PLL and Γ-distributed rate heterogeneity among sites with 4 discrete rate categories
- Use AIC, BIC, and AICc to determine the best among the 203 models
- Conduct an ML tree search (as implemented in PLL) on the best model
- Write the final ML tree to a Newick file

Note that, the actual implementation computes the best model for 5 instead of 3 information criteria. We compute BIC and AICc scores twice to allow for using distinct sample size definitions. In the following we briefly describe the sequential and parallel implementations of the algorithm.

### Sequential Implementation

Initially, we parse and read the alignment data (either in PHYLIP or FASTA format) into a PLL data structure using the appropriate I/O routines of the PLL. Besides the alignment file name, users can also specify if they want to use empirical base frequencies or conduct a ML estimate of the base frequencies. We then build a reasonable tree on which we are going to calculate the likelihoods for all models. This is done via the randomized stepwise addition parsimony tree algorithm as implemented in the PLL. We then loop over all (pre-computed) possible models and set the substitution rate dependencies in the matrix using the appropriate PLL function according to the current model string. subsequently, we optimize all model parameters (base frequencies, rates, branch lengths, α shape parameter of the Γ model of rate heterogeneity) on the fixed parsimony tree with PLL and calculate the values for the five information criteria. We keep track of the minimum value and the corresponding model for each criterion separately for further downstream analysis. Given the set of best models (at least one and at most three) according to the criteria, we then conduct a ML tree search for each criterion and best-scoring model and write the resulting ML tree to file in Newick format.

### Parallel Implementation

The runtime of the sequential implementation is dominated by likelihood calculations in the PLL that account for approximately 90 - 95% of overall execution time. Thus, the “model optimization” and “tree search” steps can become performance bottlenecks, in particular for large datasets. To this end, we designed an MPI-based Master/Worker approach that evaluates all 203 distinct models in parallel. Note that, the likelihood evaluations for each of the 203 models are completely independent of each other.

The primary goal is to efficiently find the best-fitting model for each information criterion. As already mentioned, we need to optimize the model parameters for each of the 203 models. Thereafter, we can calculate the information criteria using the respective ML score and other relevant information like the degrees of freedom and the dataset properties, as mentioned before.

Since we only need to select the “best” model per information criterion, we can merge the criterion scores of different models by keeping only the minimum^[1]^ for each criterion. This means that, if we merge the results for two distinct model evaluations, we obtain a list of lowest criterion-score/model-index pairs. To explain the parallelization we distinguish between “local” or “global” result pair lists. A *local* result only aggregates the information for a subset of models, whereas the *global* result aggregates the information over all models.

In the sequential case, the one and only thread starts with a local result list and updates it after each model evaluation. When the last model has been processed the local result corresponds to the global result.

In the parallel case each worker process maintains its own local result list. Worker processes receive work (models) from the master process, evaluate them, and update their local result lists. When all models have been evaluated, we use MPI to collect and automatically merge^[2]^ all local results lists into the global result list at the master.

In the following, we will briefly outline the MPI communication and synchronization scheme. After asynchronously sending an initial workload to each worker, the master process uses a blocking wait for any worker to finish its model evaluation task. Once a worker has finished, the master again uses an asynchronous send to propagate the next model to be evaluated to this worker. Via the asynchronous send, the master can immediately continue distributing tasks to other workers. The worker employs a blocking receive for a model, evaluates it, updates its own local result list, and informs the master about the task completion. When all models have been distributed and evaluated, the master initiates a parallel reduction operation to obtain the global result list and terminate all MPI processes.

### Code Quality Assessment

In order to continuously monitor and improve the quality of our source code, we used several tools and methods throughout our project. We deployed both, static, as well as dynamic code analyses.

#### Static Analysis

Static analyses help to identify static programming errors such as incorrect programming language syntax or static typing errors. We compiled our code with the `gcc` compiler using all available and reasonable warning flags^[3]^. This allowed us to identify potential programming errors at an early stage. Due to its more pedantic nature (i.e., ability to detect more errors), we also used the `clang` compiler with respective flags^[4]^ periodically alongside of `gcc` to further reduce the amount of potential errors.

#### Dynamic Analysis

These analyses cover run-time issues such as memory leaks. We used `valgrind` and its sub-module `memcheck` for detecting memory-related errors in our program. By appending the `Open MPI` related `valgrind` suppression file to the tool, we excluded several MPI-specific memory management functions from the analyses which resulted in an easier-to-interpret debugging output.

## Experimental Setup

To answer our main question, whether models matter or not, we applied our tool to a large number of representative empirical datasets. For each dataset we conducted 36 independent runs with 18 distinct random number seeds and 2 base frequency configurations (empirical and ML estimate) such as to generate 8 distinct randomized step-wise addition order parsimony starting trees. In each run, we determined a total of at most 5 best-scoring models using the following criteria: AIC, AICc-S (sample size: #alignment sites), AICc-M (sample-size: #alignment sites × #taxa), BIC-S (sample size: #alignment sites), BIC-M (sample-size: #alignment sites × #taxa). In addition, in each run we conducted one tree search under the GTR+Γ model.

Thus, each out of the 36 per-dataset runs generated at most 6 ML trees, 5 using the best-scoring models under the respective criteria and one under GTR+Γ. Note that, different information criteria (like the 3 variants of AIC) tend to choose the same model and therefore typically less than 6 distinct ML trees are generated overall per run. Thus, the total number of trees inferred for each dataset is ≤ 6·36 = 216. For subsequent analyses we then selected the trees with the respective highest ML score out of the 36 runs (18 random seeds) for each criterion separately and independently.

### Empirical Test Datasets

The empirical test datasets we used are summarized in Table 1. For each dataset we list the number of taxa, number of sites, and the source from which they were obtained. ‘Original paper’ refers to the Bayesian implementation in [5], ‘Lakner *et al.*’ refers to the datasets used in [16], ‘MrBayes’ refers to benchmark datasets used for testing MrBayes [6], and ‘RAxML’ refers to datasets used for benchmarking `RaxML` [9].

**Table 1.** Summary of empirical test datasets

| #taxa | #sites | source |
| --- | --- | --- |
| 46 | 1104 | Original paper |
| 64 | 1620 | Original paper |
| 13 | 1255 | Original paper |
| 106 | 378 | Original paper |
| 68 | 818 | Original paper |
| 15 | 379 | Original paper |
| 13 | 273 | Original paper |
| 23 | 2841 | Original paper |
| 17 | 379 | Original paper |
| 30 | 1456 | Original paper |
| 23 | 9741 | Original paper |
| 18 | 1689 | Original paper |
| 17 | 432 | Original paper |
| 31 | 1140 | Original paper |
| 12 | 894 | Original paper |
| 27 | 1949 | Lakner et al. |
| 29 | 2520 | Lakner et al. |
| 36 | 1812 | Lakner et al. |
| 41 | 1137 | Lakner et al. |
| 43 | 1660 | Lakner et al. |
| 50 | 1133 | Lakner et al. |
| 59 | 1824 | Lakner et al. |
| 64 | 1008 | Lakner et al. |
| 71 | 1082 | Lakner et al. |
| 50 | 378 | Lakner et al. |
| 67 | 955 | Lakner et al. |
| 4 | 16119 | MrBayes |
| 12 | 898 | MrBayes |
| 9 | 720 | MrBayes |
| 123 | 1606 | MrBayes |
| 150 | 1269 | RAxML |
| 218 | 2294 | RAxML |
| 354 | 460 | RAxML |
| 404 | 13158 | RAxML |
| 500 | 1398 | RAxML |
| 10 | 705 | RAxML |
| 101 | 1858 | RAxML |
| 714 | 1241 | RAxML |

### Hardware Platform

With increasing number of taxa, the computations can become time-consuming. Thus, we used a comparatively large multi-core shared memory machine for our experiments (running Ubuntu 14.04). The server comprises four AMD Opteron 6174 processors, each equipped with 12 cores running at 2.2 GHz and has 256 GB RAM. While our parallel implementation can use all 48 cores simultaneously, the computations for all test datasets still required several days.

## Experimental Results

In the following we describe the results of our experiments. Initially, we verify that our model generation algorithm is correct. Then, we provide speedup values for the parallel implementation. After these technical issues, we assess the magnitude of topological differences between trees inferred under GTR+Γ and trees inferred under the best-fit model according to the respective information criterion. Finally, we quantify differences between AICc, BIC, and AIC and also assess if the sample size definition has an impact on model selection.

### Verification of Model Generation Algorithms

We verified the correctness of the two model generation algorithms by systematically comparing the respective outputs to each other. Fortunately, they yield exactly identical sets of model strings. Note that, the two algorithms were developed and implemented completely independently. In addition, we also verified that our set of model strings was identical to those provided in [5].

### Speedups

To assess the parallel efficiency of our code, we measured the speedups for an increasing number of MPI processes. We only measured speedups for the model evaluation phase and excluded the tree search phase which is calculated by a stand-alone PLL function. The obtained execution time over # cores plots are provided in Figure 2 and compared to a linear speedup line. Based on our results, running the code with 6, 12, or 24 cores shows a higher parallel efficiency than with 48 cores. Nonetheless, with respect to time-to-completion, executing the code with 203 MPI processes for the 203 models will yield the fastest response.

**Figure 2.**
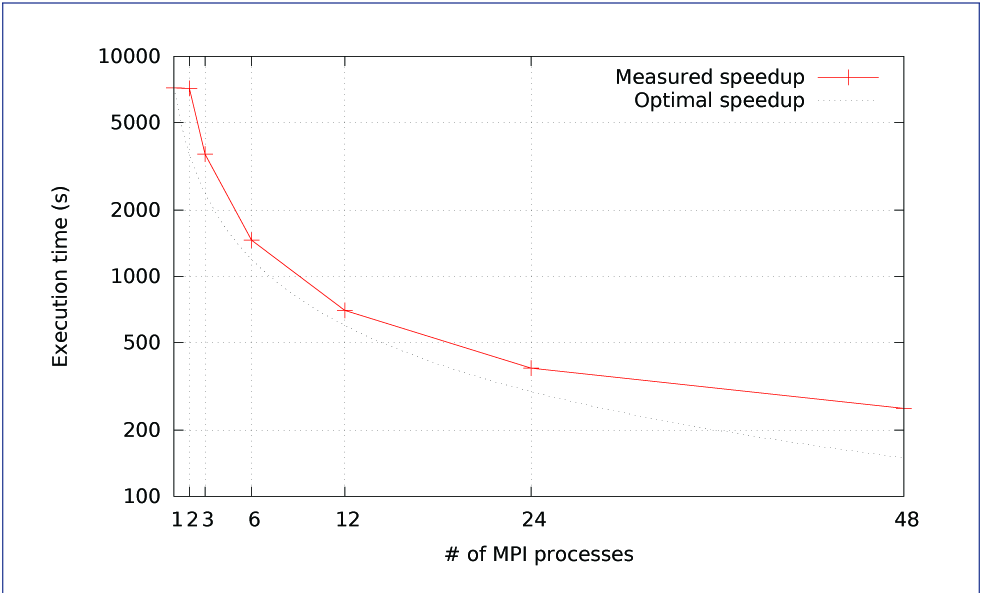
Speedup of model evaluation. Comparison of the theoretical and practical speedup of our application.

### Topological Analysis (RF Distances)

Initially, we asked the question if model selection has an impact on the final tree topology with respect to trees inferred under GTR+Γ. To this end, we calculated the relative Robinson-Foulds (rRF [17]) distances (using RAxML) between the best-scoring ML tree under GTR+Γ and the respective best-scoring trees under the 5 criteria. In Figures 3, 4, 5, 6, and 7 we show histograms of the distribution of rRF distances between the tree inferred under GTR+Γ and the trees inferred under AIC, AICc-S, AICc-M, BIC-S, and BIC-M respectively. Note the log scale on the y-axis (fraction of test datasets) of the histograms.

While for the vast majority of datasets, the rRF distance between the tree inferred under GTR+Γ and the tree inferred under the respective optimal model is zero, there is a substantial number of outliers. For more than 5% of the tree inferences the rRF distance is larger than 10% which represents a notable topological difference.

**Figure 3.**
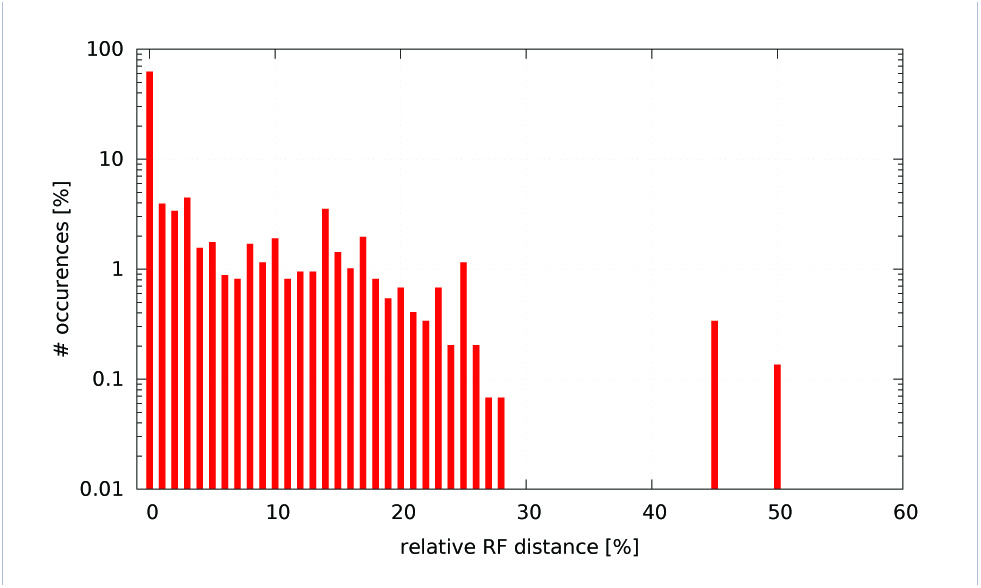
GTR+Γ model - AIC. Histogram of occurences of relative RF distances between the GTR+Γ model and the best model according to AIC.

**Figure 4.**
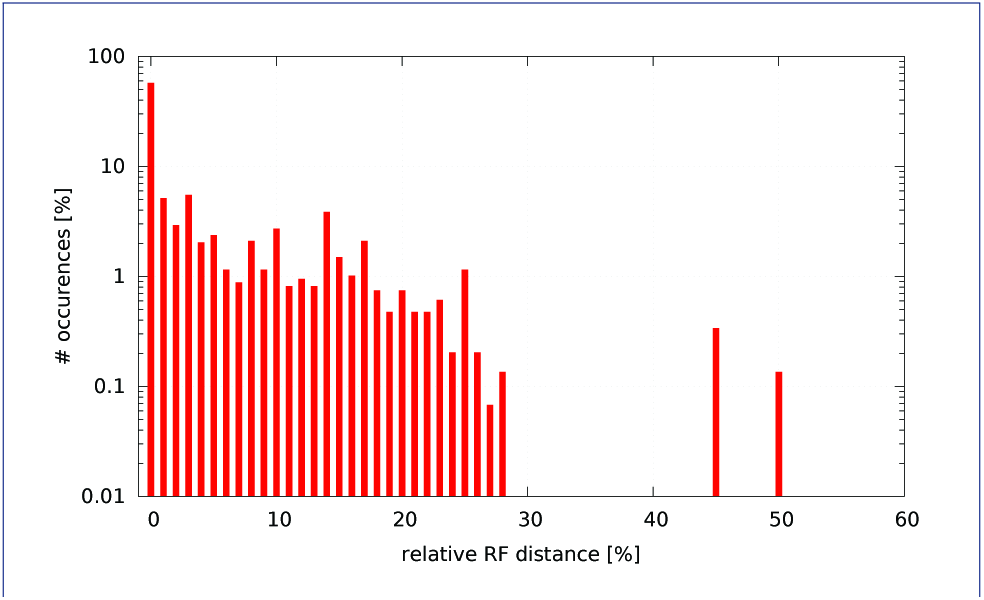
GTR+Γ model - AICc-S. Histogram of occurences of relative RF distances between the GTR+Γ model and the best model according to AICc-S.

**Figure 5.**
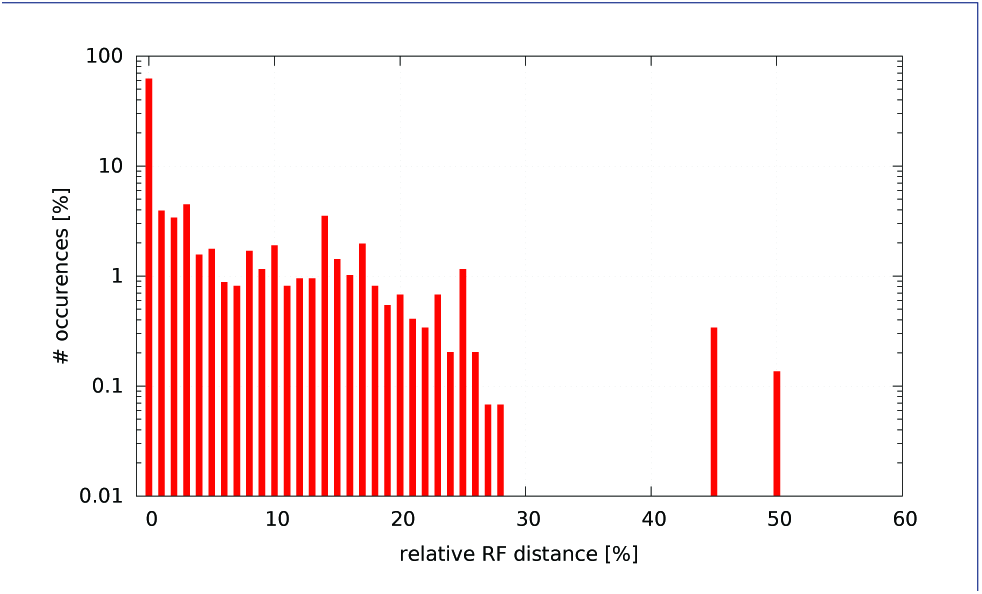
GTR+Γ model - AICc-M. Histogram of occurences of relative RF distances between the GTR+Γ model and the best model according to AICc-M.

**Figure 6.**
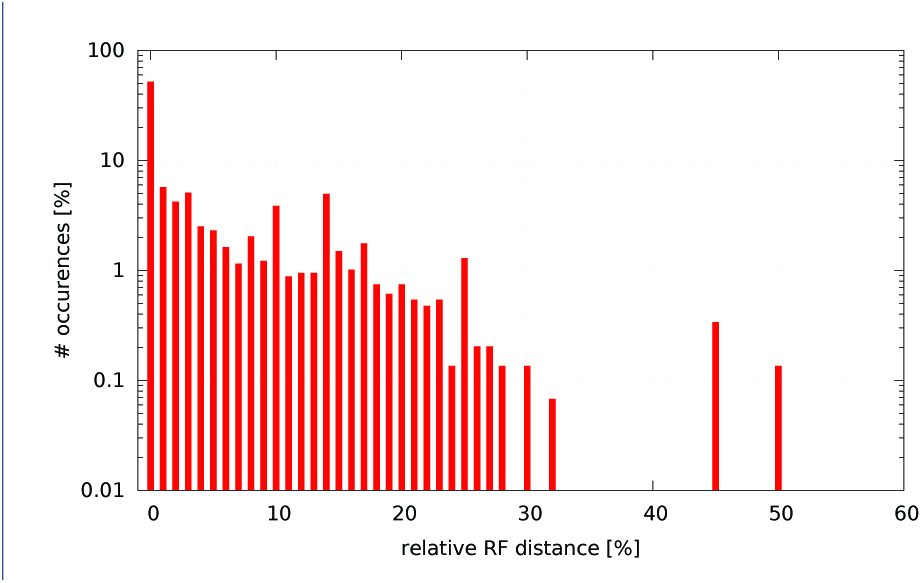
GTR+Γ model - BIC-S. Histogram of occurences of relative RF distances between the GTR+Γ model and the best model according to BIC-S.

**Figure 7.**
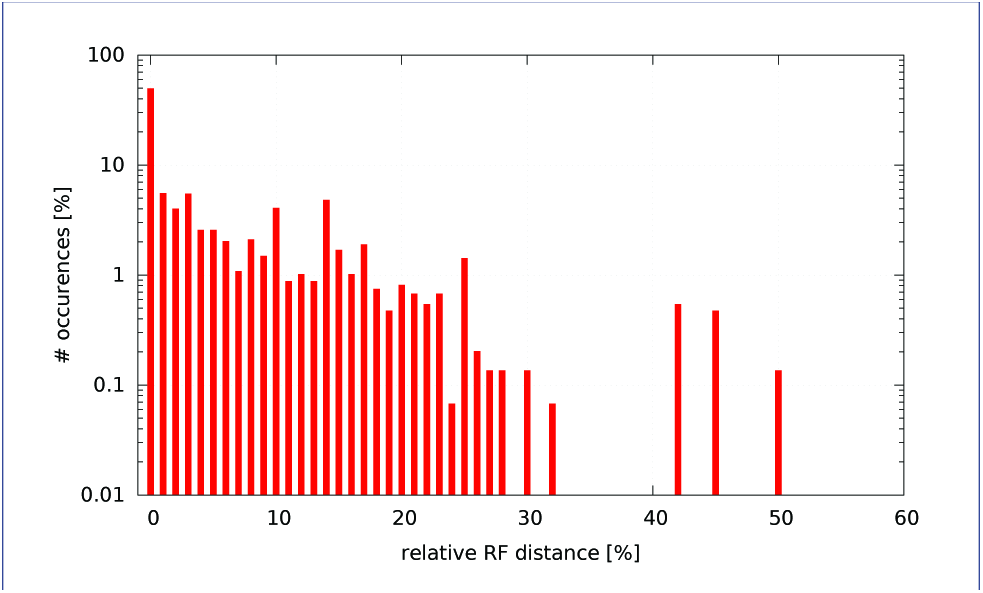
GTR+Γ model - BIC-M. Histogram of occurences of relative RF distances between the GTR+Γ model and the best model according to BIC-M.

### Differences between AICc, BIC, AIC

In the following, we quantify the differences in tree topologies and also in inferred models between the different criteria (AIC, AICc, BIC). To simplify the presentation, we only use AICc-S and BIC-S, since the number of sites is commonly used as sample size.

For comparing the inferred models, we used the model that yielded the best-scoring ML tree under AIC, AICc-S, and BIC-S, respectively. We found that AIC and AICc-S propose the same model for 1181 out of 1368 dataset configurations (38 datasets times 36 independent runs per dataset). Further, AIC and BIC-S select the same model in 683 out of 1368 cases. Finally, AICc-S and BIC-S suggest the same model for 731 out of 1368 dataset configurations.

For the cases where the proposed models between the criterion pairs (i.e., AIC/AICc-S, AIC/BIC-S, AICc-S/BIC-S) are different, we then also quantified the differences among tree topologies. For this purpose we provide histograms (Figures 8, 9 and 10) of the pair-wise RF-distances between the best-scoring trees for the three aforementioned criterion pairs (note the log scale on the y-axis).

**Figure 8.**
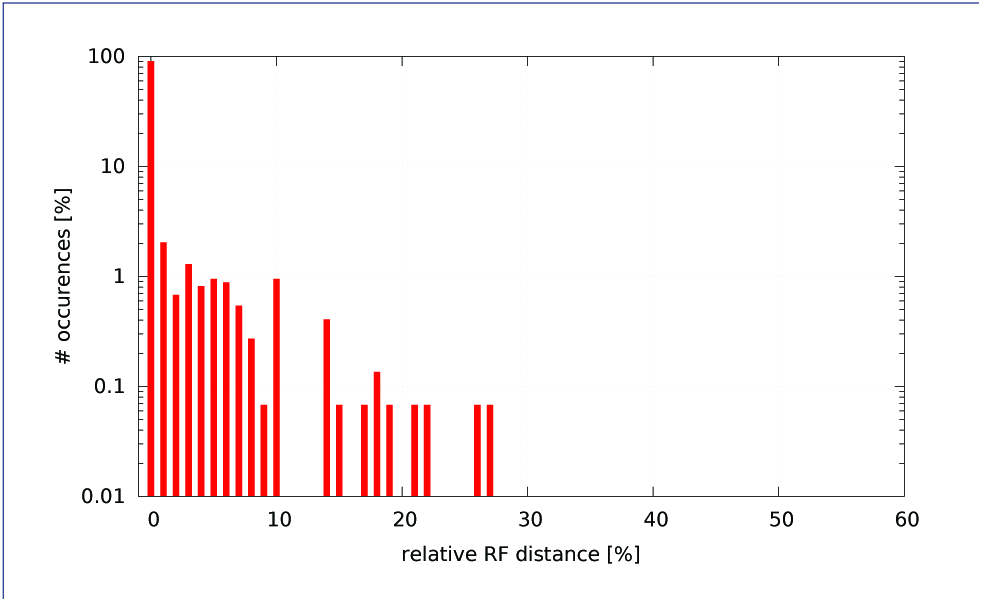
AIC - AICc-S. Histogram of occurences of relative RF distances between the best model according to AIC and the best model according to AICc-S.

**Figure 9.**
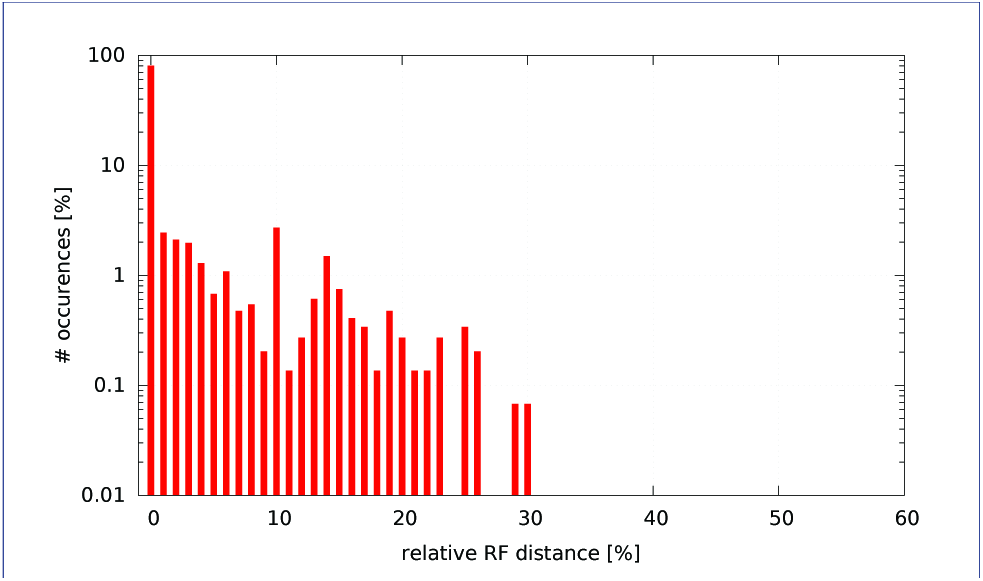
AIC - BIC-S. Histogram of occurences of relative RF distances between the best model according to AIC and the best model according to BIC-S.

**Figure 10.**
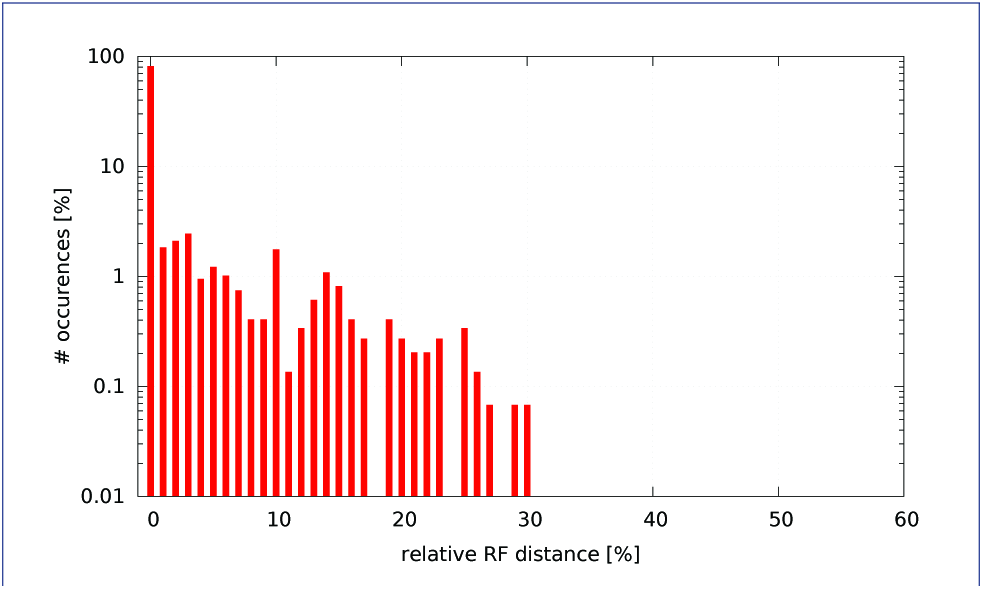
AICc-S - BIC-S. Histogram of occurences of relative RF distances between the best model according to AICc-S and the best model according to BIC-S.

We observe that, in cases where the proposed models differ, this difference can induce notable rRF distances among the inferred trees.

#### Impact of Sample Size Definition

In these analyses, we determined the impact of the sample size definition on the models selected by the AICc and BIC criteria. When comparing the model inferred for the respective best-scoring ML tree for AICc-S versus AICc-M, we found that they yielded different models for 187 out of 1368 datasets. Analogously, for BIC-S versus BIC-M different models were proposed in 261 out of 1368 cases.

As before, we then calculated the rRF distances between trees inferred under models proposed by AICc-S/AICc-M and BIC-S/BIC-M, but again only for those datasets where the models differed. The respective rRF distance histograms are provided in Figures 11 and 12 (note the log scale on the y-axis).

**Figure 11.**
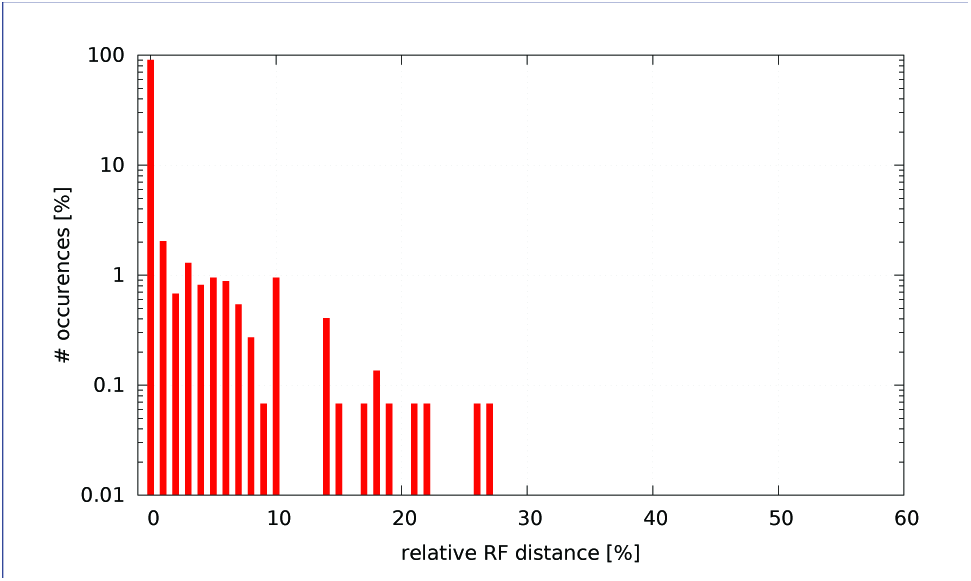
AICc-S - AICc-M. Histogram of occurences of relative RF distances between the best model according to AICc-S and the best model according to AICc-M.

**Figure 12.**
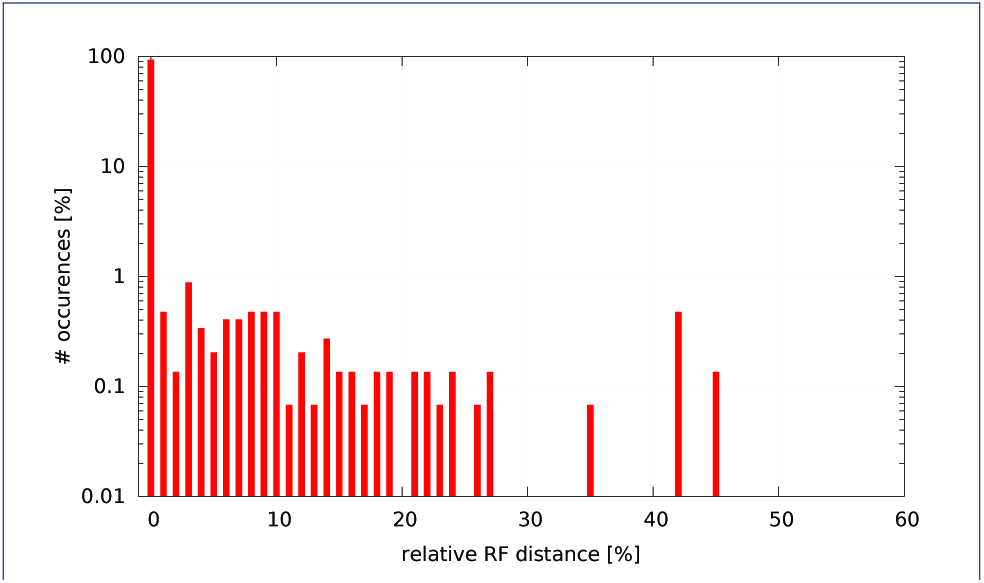
BIC-S - BIC-M. Histogram of occurences of relative RF distances between the best model according to BIC-S and the best model according to BIC-M.

As in previous experiments, there are a few dataset where the definition of the sample size yields trees that exhibit large topological distances exceeding 10%.

#### Selected Models

Another noteworthy observation is that, regardless of the criterion deployed, the subset of models that were finally selected (only 37 out of 203 possible models) is comparatively small. The type and frequency of models selected by each criterion is depicted in Figure 13.

**Figure 13.**
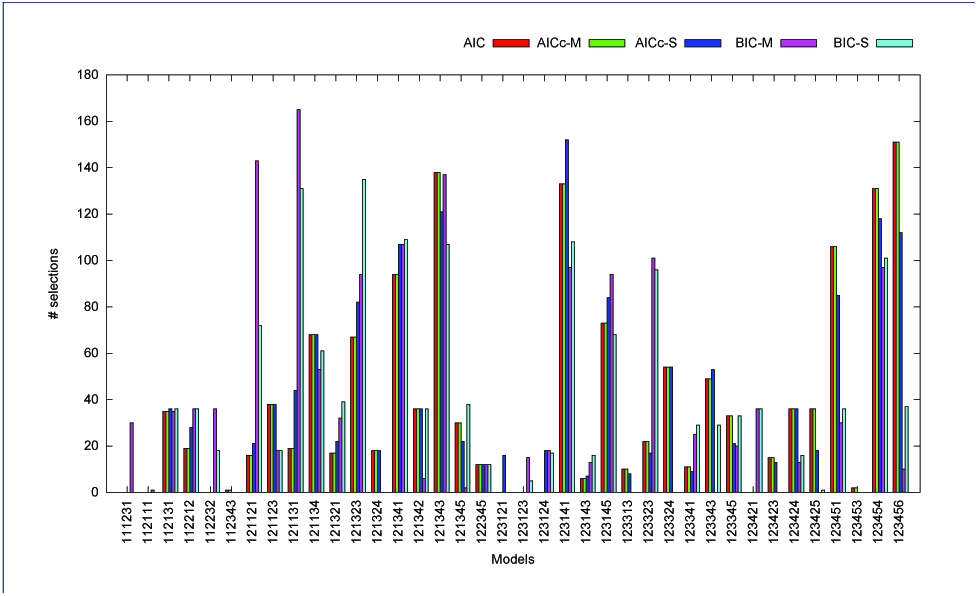
Actual subset of models selected by the distinct information criteria. The histogram shows the number and type of models selected by each information criterion for all datasets. Note that, the vast majority of possible time-reversible models was never selected.

This observation could be used for accelerating model selection by only using this subset of 37 promising candidate models.

### Significance of Results

To assess the significance of our results we conducted 50 distinct ML searches under the respective best-fit model and 50 ML searches under the GTR+Γ model for each dataset. We then calculated the pair-wise rRF distances between all pairs of trees from the first (best-fit model) and second (GTR+Γ) tree set. Thereafter, we compared the average rRF distance among these tree pairs with the random expectation for each empirical dataset. The random expectation is defined as the average rRF distance among all pairs of the 50 trees obtained from the searches under GTR+Γ. We observed significantly higher average rRF distances than the random expectation between the trees inferred under the best-fit model and GTR+Γ in 14 out of 41 datasets (p < 0.01). Inversely, for 16 out of 41 datasets, we observed differences that were significantly lower than the random expectation (*p* < 0.01). For the remaining 11 datasets, the inferred ML topology was identical, regardless of the model and the tree search replicate.

### Model Selection Accuracy under Simulation

In order to further evaluate the accuracy of our model selection process, we created 100 synthetic alignments based on the two empirical datasets (alignment with 27 taxa and 1949 sites from [16], and lice dataset from [5]) which exhibit the highest topological variance. To simulate, we initially determined the best-fit model, best-known ML tree, and optimized the model parameters for the two datasets. We then deployed INDELible [18] to generate simulated alignments.

Table 2 summarizes the results of our simulations. For single tree searches under the respective best fit model according to our model selection criteria and under GTR+Γ we quantify accuracy by showing average values (over 100 simulation replicates) for (i) recovery rate of the true topology, (ii) RF distance to the true topology, (iii) branch score (BS) difference to the true tree, and (iv) recovery rate of true, generating model.

**Table 2.** Accuracy of model selection for 100 simulations based on two ‘difficult’ empirical datasets. TTR = fraction of recovery of true topology, RF = average relative Robinson-Foulds distance to the true tree over 100 simulation replicates, BS = average branch score difference to the true tree over 100 simulation replicates, TMR = fraction of true, generating model recovery.

|  |  | TTR | RF | BS | TMR |
| --- | --- | --- | --- | --- | --- |
| Dataset 1 | AIC | 0.10 | 0.176 | 0.057 | 0.34 |
|  | AICc-S | 0.09 | 0.179 | 0.057 | 0.40 |
|  | AICc-M | 0.10 | 0.176 | 0.057 | 0.34 |
|  | BIC-S | 0.09 | 0.176 | 0.055 | 0.51 |
|  | BIC-M | 0.10 | 0.174 | 0.054 | 0.40 |
| | GTR+$\Gamma$ | 0.09 | 0.173 | 0.059 | 0.00 |
| Dataset 2 | AIC | 0.35 | 0.036 | 0.001 | 0.49 |
|  | AICc-S | 0.35 | 0.036 | 0.001 | 0.48 |
|  | AICc-M | 0.35 | 0.036 | 0.001 | 0.49 |
|  | BIC-S | 0.35 | 0.036 | 0.001 | 0.14 |
|  | BIC-M | 0.35 | 0.036 | 0.001 | 0.02 |
| | GTR+$\Gamma$ | 0.35 | 0.037 | 0.001 | 0.00 |

The perfect topology was recovered more slightly more frequently with selection criteria using the sequence length times the number of taxa as sample size (0.10 vs. 0.09). Inferences conducted under model determined by BIC show lower BS distances than the other criteria or GTR+Γ (0.54 – 0.55 vs. 0.57 – 0.59). However, these differences are not significant given the number of simulation replicates. We also observed that, when using only the sequence length as sample size, the true generating model was recovered more frequently (0.51 vs. 0.34 – 0.40).

For the second dataset (*lice*), we did not observe notable differences in any of our topological accuracy measures neither under the models selected by our criteria nor under GTR+Γ.

All selected best-fit models as well as the GTR+Γ model recovered the true topology with the same frequency (0.35). However, AIC performed much better than BIC in recovering the true generating model (0.48 – 0.49 vs. 0.02 – 0.14). For this dataset though, the recovery of the true model does not affect topolog-ical accuracy.

## Discussion & Conclusions

We implemented, parallelized, and make available an efficient tool for determining the best nucleotide substitution model among all 203 possible models using the AIC, AICc, and BIC criteria. Apart from achieving our teaching goals, and beyond the computational speed of the tool, we addressed the question if nucleotide model selection matters topologically. To the best of our knowledge, this question has not been addressed using such a large collection of empirical datasets and taking into account all possible models before. Furthermore, we analyzed the topological differences between trees inferred under models proposed by the three standard criteria and assessed if these differences are significant. Finally, we also assessed to which extent the definition of the sample size in the criteria has an effect on the selected models and inferred tree topologies.

Overall, we find that, everything matters topologically. In particular, our experiments suggest that, selecting the best-scoring out of 203 nucleotide substitution model changes the final ML tree topology (compared to an inference under GTR+Γ) by over 10% for 5% of the tree inferences we conducted. Thus, clearly, such a model test should be executed prior to starting tree inferences.

With respect to the observed topological differences induced by using distinct information criteria and sample size definitions, the effect is less pronounced since it affects a smaller fraction of datasets. Nonetheless, in some cases we did obtain substantial topological differences. However, whether one should use AIC, AICc, and BIC and how one should define the sample size is still subject to on-going statistical debates. Thus, we are not in a position to make a clear statement here. What we can state though, is that, the statistical discussions are justified based on the differences we observed.

### Code & Data Availability

The code, test datasets, and results are available under GNU GPL at https://github.com/team-pltb.

## Author’s contributions

M.H., S.O., and B.R. (the students) implemented the tool and conducted the experiments. A.S. taught the course and designed as well as supervised the programming task. D.D. provided support, including bug fixes, for the PLL and also helped with the information criteria. All authors contributed to drafting and writing the manuscript.

## Acknowledgments

We wish to thank John Huelsenbeck for providing the test datasets from the original paper, Andre J. Aberer for providing additional test datasets, and Lucas Czech for implementing an initial prototype version for the task to verify if it was ‘doable’.

## Footnotes

[1] A lower score indicates a better model

[2] intercommunicator reduce operation

[3] -Wall-Wextra-Wredundant-decls-Wswitch-default-Wimport-Wno-int-to-pointer-cast-Wbad-function-cast-Wmissing-declarations-Wmissing-prototypes-Wnested-externs-Wstrict-prototypes-Wformat-nonliteral-Wundef

[4] -Weverything-pedantic

